# dentist: Computing uncertainty by sampling points around maximum likelihood estimates

**DOI:** 10.1101/2023.01.10.523430

**Authors:** James D. Boyko, Brian C. O’Meara

## Abstract

1. It is standard statistical practice to provide measures of uncertainty around parameter estimates. Unfortunately, this very basic and necessary enterprise is often absent in macroevolutionary studies.
2. dentist is an R package allows an estimate of confidence intervals around parameter estimates without an analytic solution to likelihood equations or an approximation based on local curvature at a peak. This package works by “denting” the likelihood surface by sampling points a specified distance around the maximum likelihood estimate following what is essentially a Metropolis-Hastings walk.
3. We describe the importance of estimating uncertainty around parameter estimates as well as demonstrate the ability of dentist to accurately estimate confidence intervals.
4. We introduce several plotting tools to visualize the results of a dentist analysis. dentist is freely available from https://github.com/bomeara/dentist, written in the R language, and can be used for any given likelihood function.

## 1 Introduction

It is standard statistical practice to provide measures of uncertainty around parameter estimates. Unfortunately, this very basic and necessary enterprise is often absent in macroevolutionary studies. Problems with parameter estimation can be readily apparent empirically if one can accurately assess the confidence of these estimates and the uncertainty around the best estimate. Instead, hundreds of papers reported only point estimates (Louca and Pennell 2020).

Confidence intervals (CI) are defined by their upper and lower bounds. These endpoints represent the range of possible values which are reasonably plausible estimates for a fitted model. For example, in studying diversification rates we may find that the most likely estimate of the extinction rate of a clade is 0 in an empirical phylogeny, but when confidence intervals are examined, the range of plausible values includes values much greater than 0 (and perhaps values lower than 0, see Louca and Pennell 2020). Importantly, the biological interpretation of an extinction rate of *certainly 0* will differ massively than if it were treated as *possibly 0*. The latter interpretation highlights the difficulty in estimating a particular parameter, whereas the former can easily lead to an overinterpretation of results. Another important aspect of confidence intervals for phylogenetic comparative models is that they are often asymmetric. Although it is mathematically possible for extinction to be estimated below 0 (see Louca and Pennell 2020), the biological meaning of such an estimate is nonsensical. For birth-death models the lower confidence interval is bounded at 0, but the upper confidence interval can include any non-negative number. Issues can arise in cases such as this when confidence intervals assume a normal and symmetric distribution of error.

Several approaches exist for estimating confidence in parameter estimates. For some basic distributions, such as the binomial, a closed form solution to standard errors exists from general likelihood theory and thus we can directly solve for the sampling variance around the estimator. However, for more complex models, such as those commonly used in phylogenetic comparative methods, there are no closed form solutions for confidence intervals. A second option would be to examine the curvature at the peak of the likelihood surface to quantify the information contained in the dataset. Under assumptions of normality, the curvature can be used to derive confidence intervals (Edwards 1984). However, when those assumptions are broken (e.g., non-linearity, non-symmetry of error) this method will give incorrect estimates of the uncertainty (Wieland et al. 2021). The parametric bootstrap is able to produce confidence intervals for parameters without closed form solutions to maximum likelihood equations at the cost of increased computational effort (Efron 1987). This useful method has been applied in comparative methods to some degree (Felsenstein 1985; Boettiger et al. 2012; Jhwueng 2013), but perhaps owing to the computational effort required has not seen wide application. In cases where the true model is much more complex than the fitted model (essentially always true) simulated datasets may also be less messy than the empirical dataset: simulating under a normal distribution will produce estimates clustered around the peak, hiding any problems from the empirical likelihood surface being nearly bimodal, for example. Bayesian approaches will also naturally lead to an estimate of uncertainty if the underlying Markov chain runs well (Carlin and Chib 1995). Even weakly informative priors also could tend to obscure cases where the data are providing no insight into the fit (Alfaro and Holder 2006). Finally, using profile likelihoods one could examine univariate uncertainty by holding all parameters but one parameter at their maximum likelihood estimate (MLE) and varying the focal parameter (Venzon and Moolgavkar 1988; Meyer and Hill 1992; Murphy and Van Der Vaart 2000). However, this approach may be problematic if there is a ridge in multivariate parameter space which would not be noticed when considering parameters univariately.

The ideal way to assess the joint confidence intervals of several parameters would be to examine every possible value of the parameters. Unfortunately, this is usually not a viable option due to computational limits, so approximations are necessary. Here we introduce the R package Dentist (R Core Team 2022). Dentist works by denting the likelihood surface in order to assess confidence around a particular point estimate by examining the shape of the multivariate likelihood surface. This allows us to detect ridges in the likelihood surface which may be hidden by the assumptions of analytic approaches or may be missed from the computational limitations of other approaches. We apply dentist algorithms to several phylogenetic comparative R packages and demonstrate how it can be used for the generation of confidence intervals for any set of parameters for which a likelihood function is available. To that end, we discuss the summary and plotting functions within Dentist as well as how to interpret “good” or “bad” parameter estimates.

## 2 Theory and usage

### 2.1 The underlying algorithm

The basic procedure of Dentist is to sample points a specified distance around the maximum likelihood estimate. Dentist is initialized with a vector of parameters provided by the user which are typically the maximum likelihood estimate. New parameter values are then proposed by sampling a normal distribution with a mean equal to a vector of the original parameter values and standard deviation specified by the user. The standard deviation can be constant or set to differ for each parameter. Each sample may then “dent” the likelihood surface if the sampled point is better than the original negative log likelihood (-LnLik) plus user-set bias (2 log likelihood units by default). We note that Dentist uses the-LnLik so that successful “dents” are greater than the original-LnLik (typically the optimal value) but less than the user bias plus the optimal negative log likelihood. Points are sampled around the new rim following what is essentially a Metropolis-Hastings walk. As Dentist walks around the rim, it adjusts the proposal width so that it samples points around the desired likelihood. It does this by “tuning” the proposal width. After n steps (n `= adjust_width_interval =` 100, by default), Dentist will evaluate whether it is moving too far away from the desired likelihood or if it is staying in areas better than the likelihood of the desired ridge. If more than 30% of the most recent n steps resulted in an accepted “denting” of the likelihood surface, proposal widths are increased by 50% (1.5 * `sd_vector`). In contrast, if less than 10% of the most recent n steps resulted in an accepted “denting” of the likelihood surface, proposal widths are decreased by 20% (0.8 * `sd_vector`). Finally, Dentist will expand the proposal width for parameters if extreme values still appear good enough to try to find out the full range for these values. Tuning can be adjusted by the user; the defaults often work in practice but there is no guarantee.

### 2.2 Convergence on true CIs

To test Dentist, we estimate the confidence intervals for data generated using the log normal distribution. This distribution is chosen so that the confidence intervals generated by Dentist can be compared directly to confidence intervals which are exactly defined. The 95% confidence intervals for the mean estimate of a log-normal distribution is given by 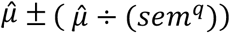 where 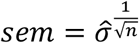 and *q* is the 97.5% quantile of the t-distribution with *n* − 1 degrees of freedom. For each iteration of our simulation procedure, we generate 100 data points with a *μ* = 1 and *σ* = 3 using `stats::rlnorm` and optimize `stats::dlnorm` to find the MLE for *μ* and *σ*. The dentist algorithm is applied with `delta=1.92` and, to better illustrate the importance of sampling many points, `nsteps` is varied to be 10, 50, 100, 500, 1000, and 5000. This procedure is repeated for 100 iterations. The accuracy of Dentist is measured as the average absolute distance between the confidence intervals generated by the closed form solution and those generated by Dentist.

We find that Dentist rapidly converges to the closed form confidence intervals as the number of steps taken increases (Fig. 1). For this example, we only estimate the confidence intervals around 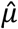, because, to our knowledge, a closed form solution for confidence intervals around 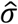 does not exist. However, Dentist in addition to estimating CIs around the mean, did estimate CIs around 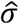. Although Dentist converged to the true CIs after between 500 and 1000 steps, if the number of freely estimated parameters increased, the algorithm would likely require more steps to accurately assess the CIs.

**Figure 1.**
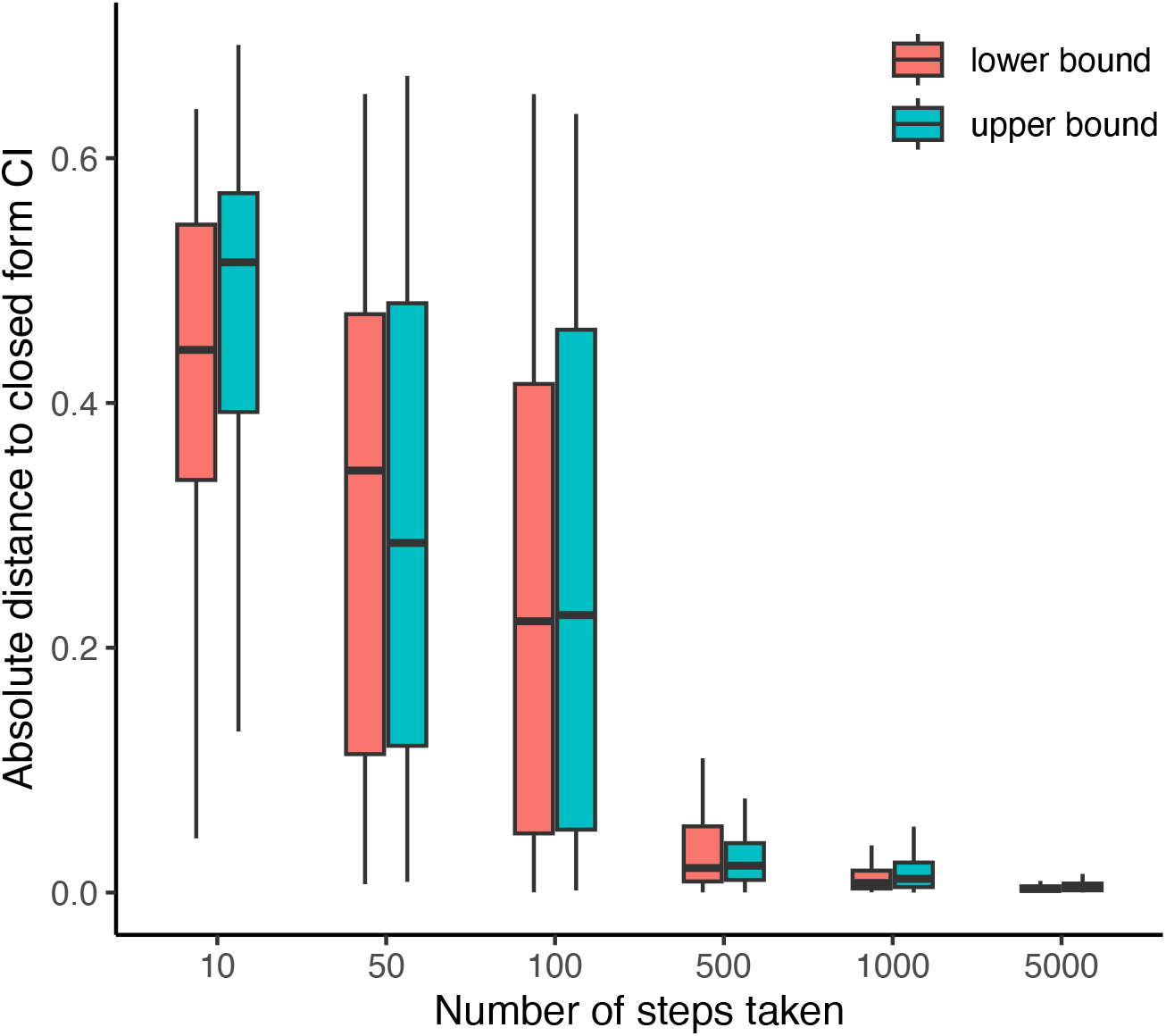
The absolute distance between the true confidence intervals and the confidence intervals generated by denting the likelihood surface. As the number of steps increases, the distance between the true confidence intervals and those generated by Dentist decreases.

### 2.3 Using Dentist

One advantage of Dentist is that it can be readily applied to any function which can take as inputs a fixed vector of parameters and outputs a likelihood. Although we cannot specify exactly how this will work for all R packages, we can demonstrate how a few simple functions allow for the analysis of confidence intervals when these two basic requirements are met. The main function of Dentist is `dent_walk()` and the arguments of this function alongside their descriptions are outlined in Table 1. Several of these inputs and why they are important have previously been discussed in section 2.1 and 2.2, but now we focus on inputs which are necessarily user-specified: `par, fn, best_neglnL`. We use the R package corHMM (Beaulieu et al. 2013; Boyko and Beaulieu 2021) as an example because it meets the two requirements of allowing the input of fixed parameters while outputting a likelihood. corHMM is a phylogenetic comparative method for estimating rates of evolution between discrete characters. In this case the confidence intervals will be a range of plausible evolutionary rates. However, the main function, `corHMM()`, does not follow the exact formatting necessary for Dentist. Therefore, using Dentist will require some functional coding, but we hope that the subsequent example will serve as a general guide for using the package for both users and package developers. This will feel familiar to people using tidyverse R functions with packages that do not use data as the first argument.

**Table 1.** The main input variables for dent_walk and a brief explanation for each.

| Argument | Definition |
| --- | --- |
| `par` | Starting parameter vector, generally at the optimum. If named, the vector names are used to label output parameters. |
| `fn` | The likelihood function, assumed to return negative log likelihoods |
| `best_neglnL` | The negative log likelihood at the optimum; other values will be greater than this. |
| `delta` | How far from the optimal negative log likelihood to focus samples |
| `nsteps` | How many steps to take in the analysis |
| `print_freq` | Output progress every `print_freq` steps |
| `lower_bound` | Minimum parameter values to try. One for all or a vector of the length of `par`. |
| `upper_bound` | Maximum parameter values to try. One for all or a vector of the length of `par`. |
| `adjust_width_interval` | When to try automatically adjusting proposal widths |
| `badval` | Bad negative log likelihood to return if a non-finite likelihood is returned |
| `sd_vector` | Vector of the standard deviations to use for proposals. Generated automatically if NULL |
| `restart_after` | Sometimes the search can get stuck outside the good region but still accept moves. After this many steps without being inside the good region, restart from one of the past good points |
| `debug` | If TRUE, prints out much more information during a run |
| `...` | Other arguments to `fn`. |

As an input `corHMM()` requires several arguments for data and model structure (those arguments are at minimum `phy, data, rate.cat`). In order for Dentist to evaluate the likelihood for this model, dent_walk will also require these inputs to be the same as the fitted model. However, the input in `corHMM()` we are most concerned with is the optional argument `p`, which is a vector of transition rates (the main parameter being estimated) and allows the user to calculate a likelihood with fixed parameters. Dentist requires that this “parameter argument” be the first input of fn and so we will need to construct a function using `corHMM` where this is the case.

~~~
1 fn_corHMM <- function(par, phy, data, rate.cat){
2 corhmm_fit <- corHMM(phy = phy, data = data, rate.cat = 1, p = par)
3 loglik <- corhmm_fit$loglik
4 neg_loglik <--loglik
5 return(neg_loglik)
6 }
~~~

The above function (`fn_corHMM()`) serves to transform the main corHMM function (`corHMM()`) into one that `dent_walk()` can use. Specifically, the first argument of fn_corHMM() is par and all other arguments (`phy, data, rate.cat`) were required to specify the evolutionary model and required dataset. The key transformation occurs on line 2, where the `par` argument of `fn_corHMM()` is input as the `p` argument in `corHMM()`. This simple transformation, and ensuring that a single value for the negative log likelihood is returned (lines 3 to 5), is all the required formatting for the `fn` argument of `dent_walk()`. The other necessary inputs of `dent_walk()` are par and `best_neglnL`. These will be obtained from the initial search of your fitted model and are input directly as their numeric values. For `par`, it is recommended that the vector is named for easier interpretation of results once `dent_walk` is completed.

~~~
1 par <- c(0.01689557, 0.006224654)
2 names(par) <- c(“rate_21”, “rate_12”)
3 dent_res <- dent_walk(par=par, fn=fn_corHMM, best_neglnL=21.36498, phy=phy, data=data, rate.cat=1)
~~~

While running, `dent_walk()` will display the current range of likelihoods in the desired range (by default, the best negative log likelihood + 2 negative log likelihood units) and the number of parameter values falling in that range. If things are working well, the range of values will stabilize during a search.

~~~
[1] “Done replicate 500”
[1] “CI of values (the 266 replicates within 2 neglnL of the optimum)”
       neglnL rate_21 rate_12
[1,] 21.36498 0.004073479 0.002031896
[2,] 23.36438 0.032472031 0.016600744
~~~

### 2.4 Summarization, interpretation, and plotting

Once the search has completed, the Dentist results can be summarized and interpreted. The two main functions being used for interpretation are `summary.dentist()` and `plot.dentist()`. `summary.dentist()` takes the resulting object from a `dent_walk()` and displays summary information for each parameter. This is where the confidence intervals can most easily be extracted for each parameter as `lower.CI` and `upper.CI` rows of the summary table. This can also be examined directly as an element of the Dentist result directly through `dent_res$all_ranges`.

~~~
This ran 1000 steps looking for all points within 2 negative log likelihood units of the best parameter values.
~~~

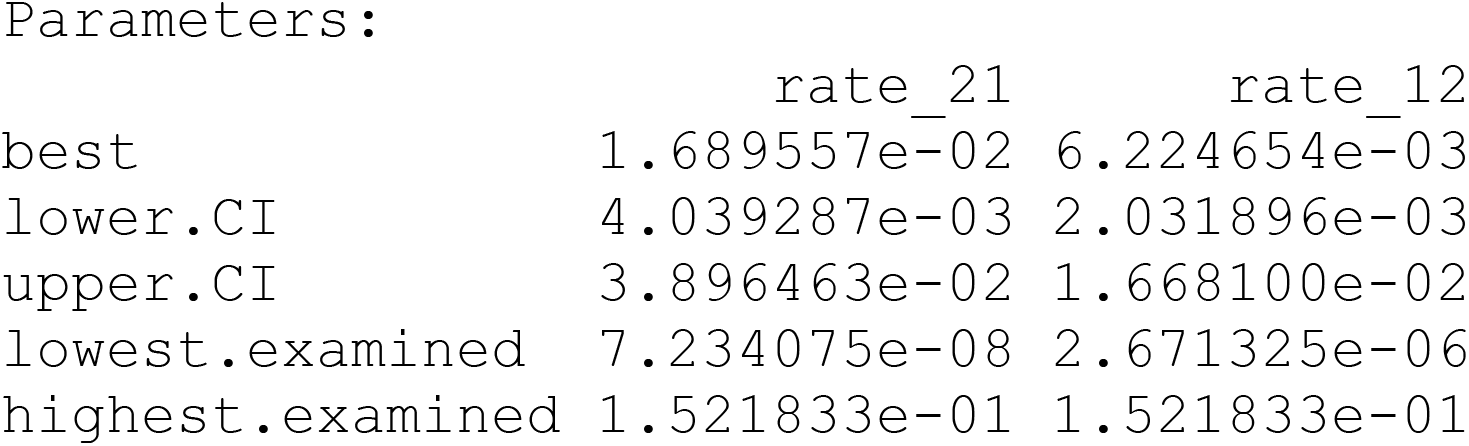

However, it can be more important to examine your model for “ridges”in parameter space that are not captured by the range of confidence intervals. This is done primarily through `plot.dentist()` which displays two types of plot (Fig. 2). First, there will be univariate plots of the parameter values versus the likelihood which can assess how well each parameter is estimated and is where the confidence intervals are derived from. The second type of plot are bivariate plots of pairs of parameters which are primarily used to look for ridges. Ridges are combinations of parameters which have likelihoods within the user range specified (`delta` argument) across parameter values. The example below shows parameters which were well estimated and do not have ridges, although there is uncertainty around the best parameter estimates. In the next section we will describe how one would detect ridges and what, if anything, can be done in those cases.

**Figure 2.**
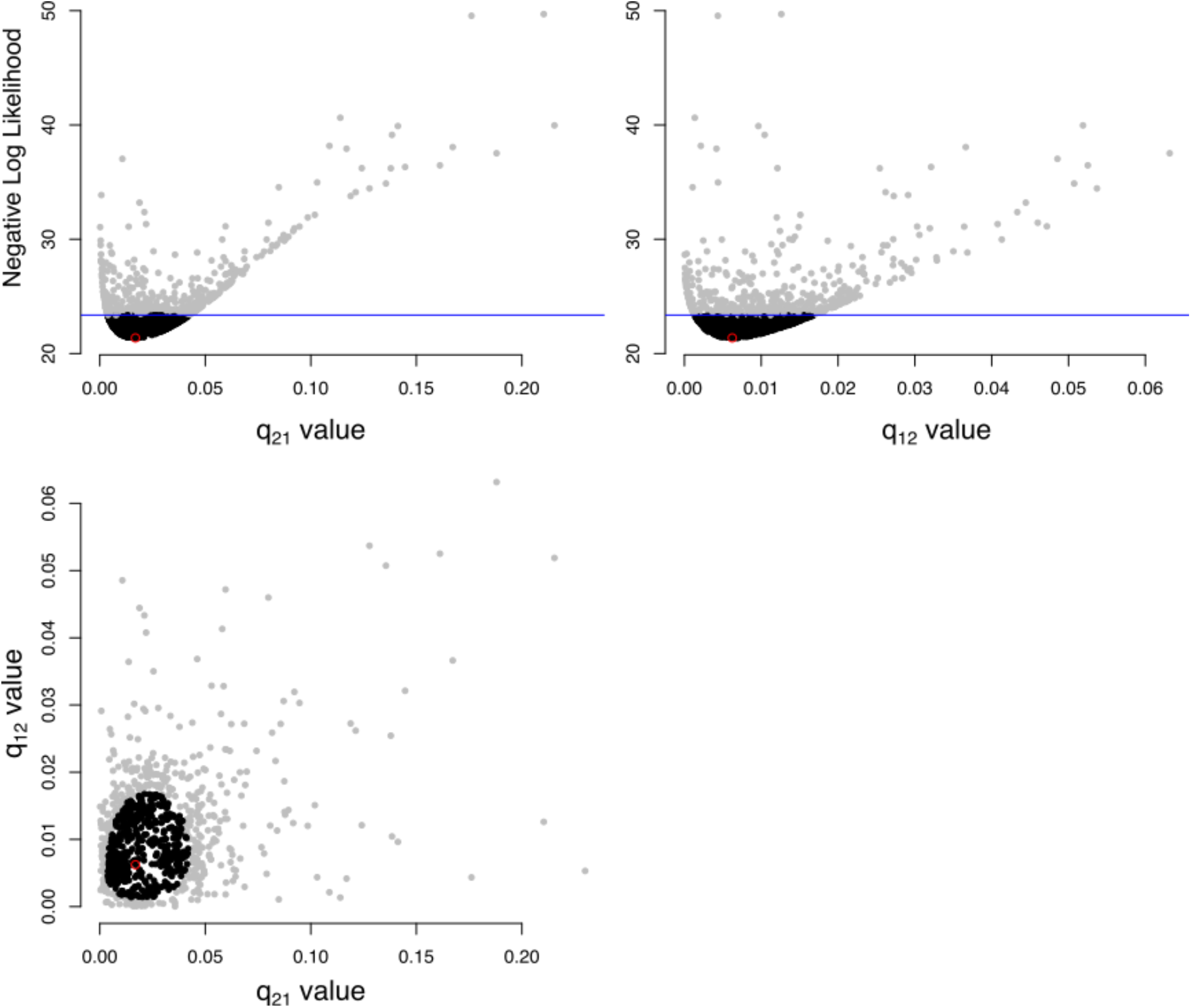
Example of plotted dentist results. There are 2 types of plots being shown here. In the first row are univariate plots displaying the negative log likelihood as a function of the parameter value. This is where the confidence intervals are shown and can assist in understanding which parameters are more or less certain than others. The remaining plot is a bivariate plot of parameters which can help look for ridges in parameter space. A ridge would appear as lines of black points. There are no ridges present in this example as all black points (within range) are clumps.

### 2.5 The good, the bad, and the ugly

A good model is one which successfully describes the data and has well-determined parameters. This is the case of the simple two-state Markov model shown in Figure 2. None of the uncertainty estimates are at their bounds, the bounds themselves are reasonably narrow, and the maximum likelihood estimate is at optimum value. In a case such as this, it is reasonable to use the model for biological insight by interpreting the parameter estimates as informing the underlying process. Ultimately, this is the goal of modeling: to gain insights that would not have been possible from examining the data alone. However, even well-defined models with theoretically estimable parameters will sometimes run into problems related to parameter uncertainty. Here we will distinguish between two major issues of parameter estimation: (1) structural unidentifiability and (2) practical unidentifiability (Wieland et al. 2021).

A common practice in macroevolution is to estimate the speciation and extinction rates from reconstructed phylogenetic trees. However, the ability to estimate extinction rates has been consistently questioned in the literature (e.g., Rabosky 2010; Louca and Pennell 2020, 2021). Nonetheless, this seeming unidentifiability of extinction rates may be a consequence of a lack of data (Beaulieu and O’Meara 2015) and therefore may instead represent an example of “practical unidentifiability” (Wieland et al. 2021). Practical unidentifiability occurs when there is not enough information in the given dataset to accurately estimate parameters and leads to a seemingly infinite number of plausible parameter values or confidence intervals at bounds. This is the case for estimating extinction rates even with simple time-homogenous birth death models (Fig. 3a). The lower bound of the extinction rate confidence intervals hits zero and the maximum likelihood estimate may even be negative in such cases (Louca and Pennell 2020). However, this seeming unidentifiability can be recovered by adding more data (Fig. 3b). Obviously, in most empirical use cases “adding more data” is not a simple solution, but that means it is all the more important to examine confidence intervals which accurately reflect the true bounds of plausible parameters to avoid making inferences with false confidence.

**Figure 3.**
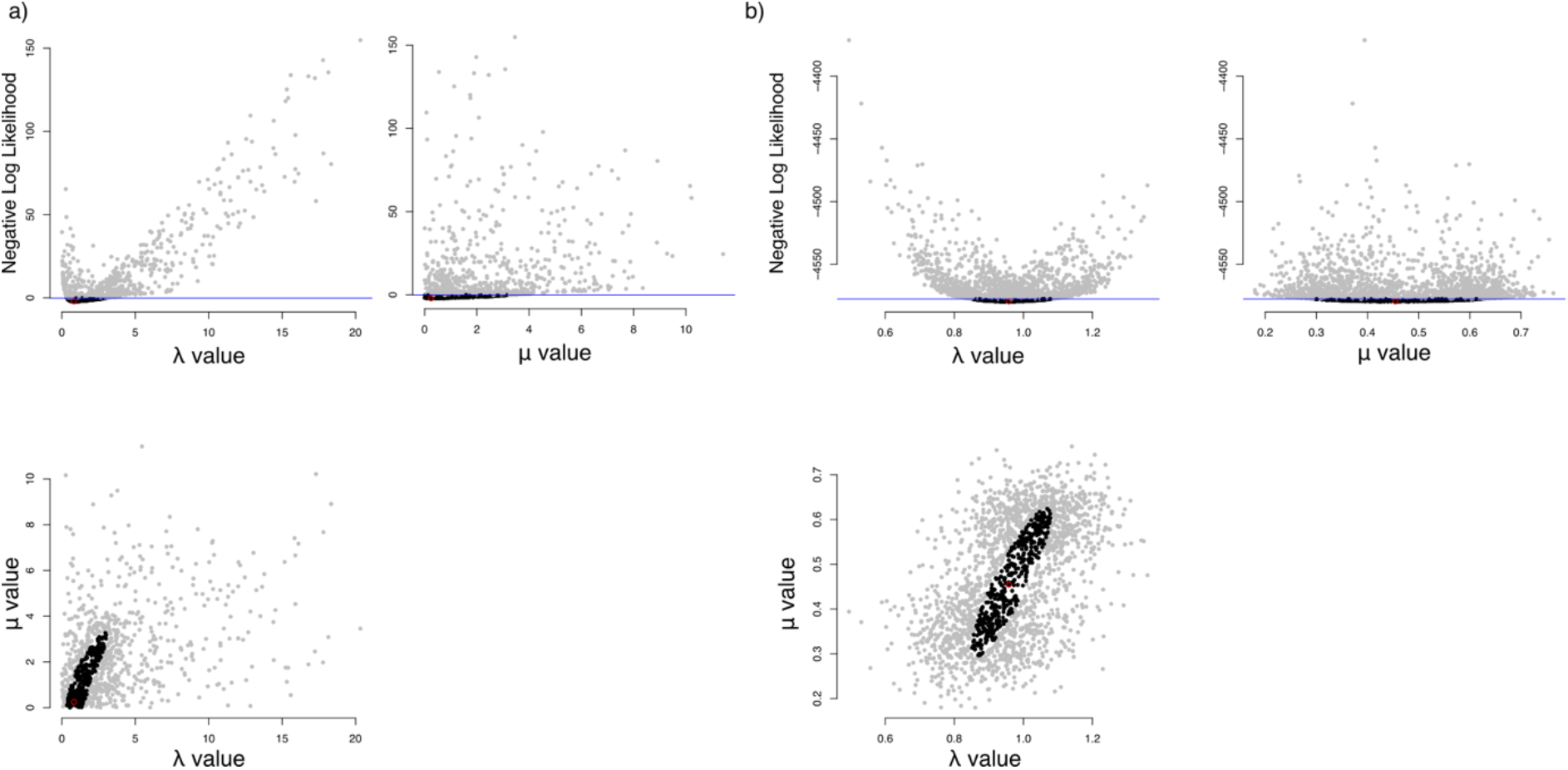
An example of practical unidentifiability resulting from a lack of data necessary to evaluate a parameter correctly. a) It is difficult to identify the extinction parameter. Mu is quite flat with a large range of possible values and its lower bound hits 0, indicating that it may not be possible to estimate this parameter correctly. b) In contrast, under the same simulation conditions, the rightward plots have identifiable parameter values although the uncertainty persists.

In contrast, structural unidentifiability cannot be addressed by additional data. In these cases, there are an infinite number of values which will result in the same likelihood. In Dentist, this will appear as a ridge in the parameter space (Fig. 4). Two simple and non-phylogenetic examples of this are the algebraic expressions *lnL* = |10 − *x* − 5*y*| (Fig. 4a) and *lnL* = 110 − *x* − *y*^2^| (Fig. 4b). The ridges appear as a set of values which will satisfy the equalities and result in confidence intervals which would theoretically extend between infinity and negative infinity. However, it is not possible to evaluate an infinite number of points and so it is left to the user to determine whether their results constitute a ridge. For these algebraic examples, we also include the posterior probabilities from a simple Bayesian Markov Chain Monte Carlo with an exponential prior. The confidence intervals produced by `dent_walk` will inevitably be finite, but the plots show ridges which *could* extend to infinity. This is in contrast to the posterior distributions which give far less of an indication that the confidence intervals of the parameter estimates could extend to infinity (Fig 4). Finally, it is exceedingly difficult to demonstrate true structural unidentifiability without the exact solution to the likelihood function, but the Dentist plots can provide partial evidence of unidentifiability.

**Figure 4.**
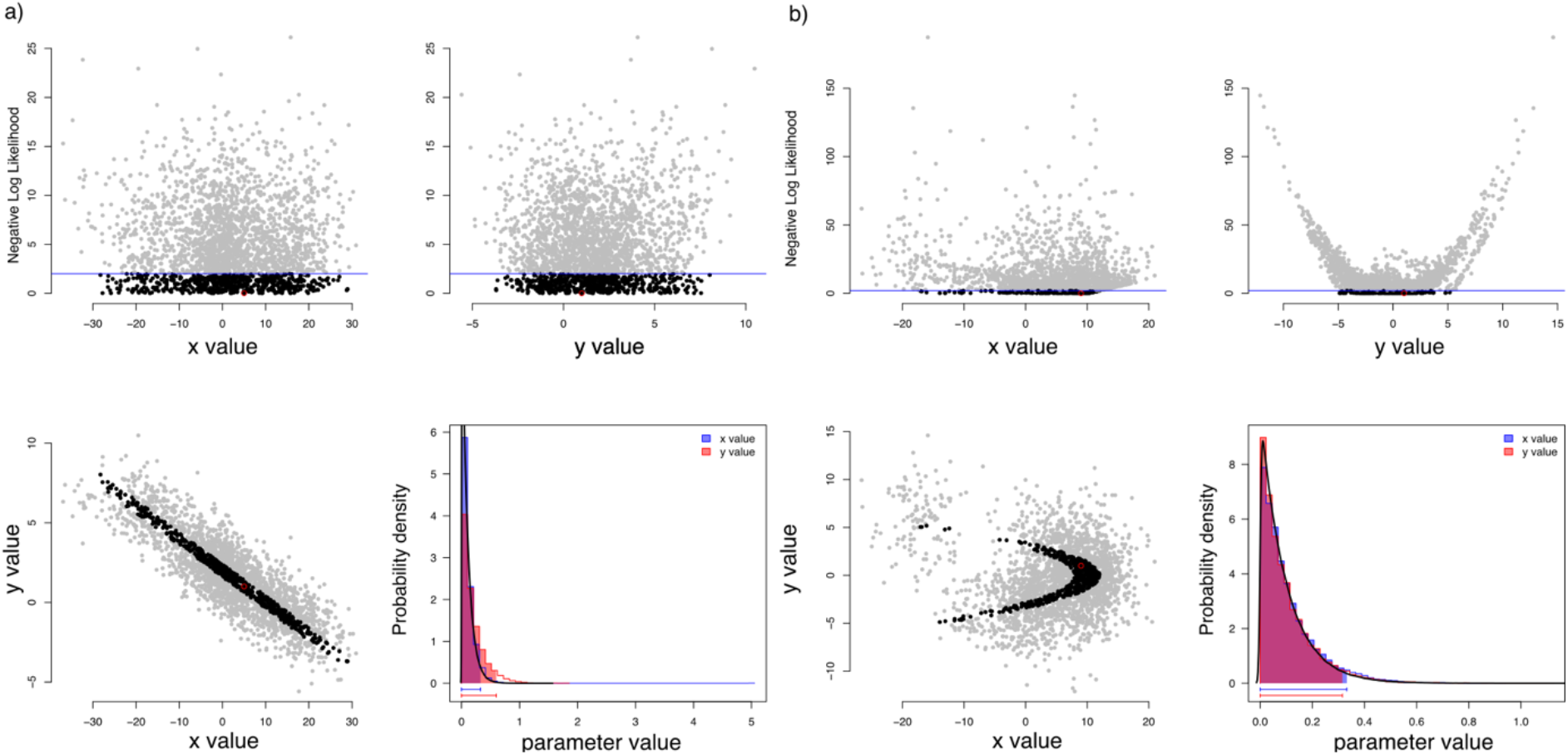
a) A simple example of fitting a model of lnL=|10-x-5y| and the resulting unidentifiable parameter estimates. A clear ridge can be seen in the bivariate y vs. x plot in which for every value of X there is a corresponding Y value which can satisfy the equality. Ridges in empirical examples will almost never be as clear as this because the models and the data are more complex. b) More complex models can result in non-linear unidentifiable parameters such as this example of lnL=|10-x-y2|. Unlike (a), the resulting ridge is not a line but a curve. Blue lines in b) represent the true CIs for this likelihood function which narrow as x approaches negative infinity. In the bottom right corner of both (a) and (b) is the posterior distributions of a simple MCMC showing finite confidence intervals. The exponential prior is overlaid via a black line and can be seen leading to seemingly finite CIs.

A more concrete example of structural unidentifiability can be found in phylogenetic comparative methods and specifically in some subsets of the Ornstein-Uhlenbeck model (Ho and Ané 2014). In this next example we fit an Ornstein-Uhlenbeck model with a variable optimum (θ), variable variance (σ^2^), and variable root value (θ_root_). This is a well-known case of unidentifiability in phylogenetic comparative models as no matter how much data is added the ridges seen in Figure 5 will persist (Ho and Ané 2014). It is simply not possible to estimate the root value under this model. However, one solution to unidentifiability is to acknowledge the informational limitations of the data and reparameterize the model itself. In the case of OU models, identifiability can be recovered by fixing the root state and this type of reparameterization is a general solution to structural unidentifiability (Wieland et al. 2021).

**Figure 5.**
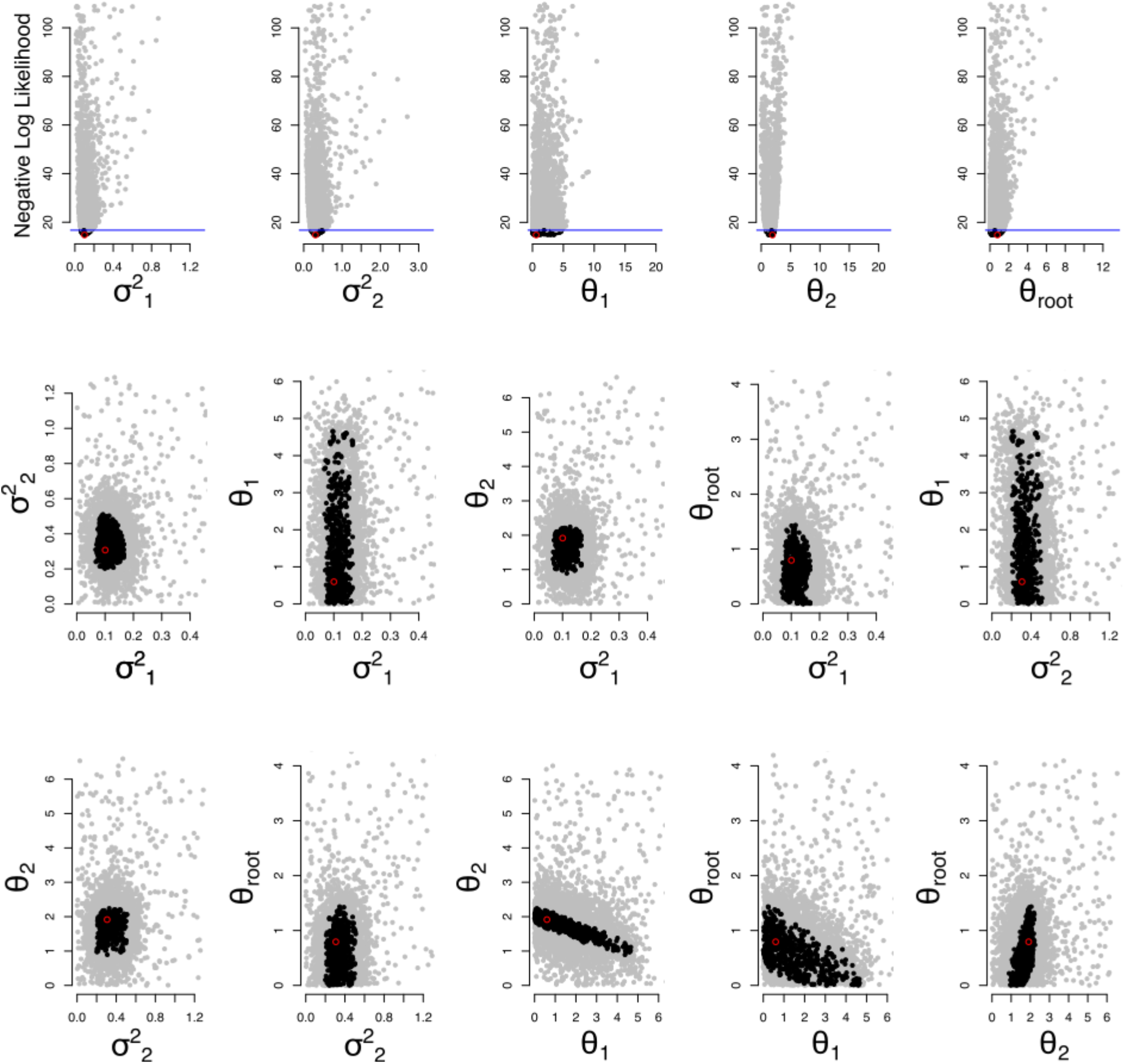
An example of unidentifiability in an OU model with a variable starting root value. Ridges can be clearly seen in several of the bivariate plots (θ_2_ vs θ_1_, θ_root_ vs θ_1_, and θ_root_ vs θ_2_) and we have highlighted those ridges with a red outline. This is also reflected in the flat uncertainty estimates in the top row of plots.

Finally, we note that both structural unidentifiability and practical unidentifiability are not necessarily the fault of the packages where the models are input. Almost any model, when coded in the most general way, can be made unidentifiable. Our work here is meant to highlight the importance of developing and using methods which have the potential to discover ridges in parameter space, detect possible unidentifiability, and reflect the accuracy of our parameter estimates through correct confidence intervals.

## 3 Discussion

Confidence intervals are an important aspect of the modeling process that can often be overlooked. In addition to an assurance of the quality of results through the quantification of plausible parameter values, they give biologists an additional tool for assessing and discussing their modeling results. The quality of modeling results is particularly important for fields like phylogenetic comparative methods, which rely on inference from observational data for insights into the evolutionary process. Issues related to statistical power and unidentifiability have been frequently acknowledged in the past, but it is necessary to remain vigilant against this problem arising in the future (Bartoszek et al. 2012; Ives and Garland 2014; Uyeda and Harmon 2014; Morlon et al. 2022). Measures of parameter uncertainty can provide insights into the identifiability or unidentifiability of particular models. However, even when confidence intervals are examined, there are limitations to both the computational and analytic approaches commonly used which can lead to overconfidence of results. Additionally, unidentifiability of models can be hidden behind the assumptions of analytic methods (Wieland et al. 2021).

The difference between structural and practical unidentifiability is an important one. Something being difficult to estimate is not the same as being impossible to estimate. Biological data is noisy and that makes statistically testing our hypotheses in observational fields such as comparative methods often difficult (Tsimring 2014). Compounding this is the fact that biologists seek to understand subtle differences that may be sensitive to practical unidentifiability if the empirical data may not contain a large signal in favor of any particular hypothesis. In the case of estimating extinction rates from phylogenies containing only extant taxa, the discussion has been framed as whether or not extinction rates can be estimated (Rabosky 2010; Louca and Pennell 2020, 2021). But, the valid estimation of extinction has been demonstrated several times and so it is known to be theoretically possible (e.g., Nee et al. 1994; Stadler 2013). In contrast, the datasets used empirically may be over parameterized for a potentially weak signal which will lead to practical unidentifiability. Nonetheless, with enough data we can have confidence in the extinction rate estimates (Beaulieu and O’Meara 2015). This is not to say that there is no structural identifiability in diversification models, there certainly are examples of this problem (Kubo and Iwasa 1995; Louca and Pennell 2020).

The final component of good statistical modeling is model selection and multimodel inference (Burnham and Anderson 2002). Often, differing model structures can be compared directly using information criteria. Under a likelihood framework, in order to compare different model structures, the most likely set of parameters of each must be found. However, the act of comparing models based on their most likely parameter estimates does not inform us about the parameter uncertainty. A model with a ΔAICc of 0.1 better than the next best model does not mean the parameters were uncertain and a ΔAICc of 20 does not mean the best fitting model had well-estimated parameters. In these cases, it remains important to accurately measure parameter uncertainty. However, this parameter uncertainty is compounded by the uncertainty in selecting a “best” model or “best set” of models (Burnham and Anderson 2002). Future work within Dentist will extend the algorithms presented here to allow for estimation of parameter uncertainty while accounting for model selection uncertainty.

## 4 Conclusion

Comparative biologists have limited power from their samples to draw inferences about macroevolutionary patterns. We cannot directly observe macroevolution and we must instead draw inferences from our observational data. Furthermore, we must always be sensitive to the limits of our models. Unidentifiability is one example of the limitations to our modeling, but as we expand the scope of study to less closely related groups, the plausibility of simple linear and homogeneous models is going to be called severely into question. Adding more realism may mean introducing additional complexity to model structure and increasing the number of parameters. As models become more complex, they are potentially in danger of having more uncertain parameter estimates and possibly being outright unidentifiable. Thus, it is important to find general ways to propagate uncertainty through our analyses and inform our inferences. Here we have shown that Dentist can augment the existing tool kit by providing reliable estimates of confidence intervals when they are finite in addition to assisting in the detection of model unidentifiability through plotting.

## Acknowledgements

We thank Jeremy Beaulieu for this thoughtful comments and feedback on earlier drafts of this work. Conversations with Tony Jhwueng led to much of the initial impetus behind the Dentist approach. This work was funded by the National Science Foundation (grant DEB-1916539)

## Author Contributions

BCO conceived the idea and coded the original package. JDB tested and updated the package. JDB and BCO wrote the manuscript.

## Data Availability

All code and simulated data are available at https://github.com/jboyko/dentist-paper.

## References

Alfaro, M. E., and M. T. Holder. 2006. The Posterior and the Prior in Bayesian Phylogenetics. Annual Review of Ecology, Evolution, and Systematics 37:19–42.

Bartoszek, K., J. Pienaar, P. Mostad, S. Andersson, and T. F. Hansen. 2012. A phylogenetic comparative method for studying multivariate adaptation. Journal of Theoretical Biology 314:204–215.

Beaulieu, J. M., and B. C. O’Meara. 2015. Extinction can be estimated from moderately sized molecular phylogenies. Evolution 69:1036–1043.

Beaulieu, J. M., B. C. O’Meara, and M. J. Donoghue. 2013. Identifying Hidden Rate Changes in the Evolution of a Binary Morphological Character: The Evolution of Plant Habit in Campanulid Angiosperms. Systematic Biology 62:725–737.

Boettiger, C., G. Coop, and P. Ralph. 2012. Is Your Phylogeny Informative? Measuring the Power of Comparative Methods. Evolution 66:2240–2251.

Boyko, J. D., and J. M. Beaulieu. 2021. Generalized hidden Markov models for phylogenetic comparative datasets. (N. Cooper, ed.) Methods in Ecology and Evolution 12:468–478.

Burnham, K. P., and D. R. Anderson. 2002. Model selection and multimodel inference: a practical information-theoretic approach (2nd ed.). Springer, New York.

Carlin, B. P., and S. Chib. 1995. Bayesian Model Choice Via Markov Chain Monte Carlo Methods. Journal of the Royal Statistical Society: Series B (Methodological) 57:473–484.

Edwards, A. W. F. 1984. Likelihood. CUP Archive.

Efron, B. 1987. Better Bootstrap Confidence Intervals. Journal of the American Statistical Association 82:171–185.

Felsenstein, J. 1985. Confidence limits on phylogenies: an approach using the bootstrap. evolution 39:783–791.

Ho, L. S. T., and C. Ané. 2014. Intrinsic inference difficulties for trait evolution with Ornstein-Uhlenbeck models. Methods in Ecology and Evolution 5:1133–1146.

Ives, A. R., and T. Garland. 2014. Phylogenetic Regression for Binary Dependent Variables. Pages 231–261 in L. Z. Garamszegi, ed. Modern Phylogenetic Comparative Methods and Their Application in Evolutionary Biology: Concepts and Practice. Springer, Berlin, Heidelberg.

Jhwueng, D.-C. 2013. Assessing the Goodness of Fit of Phylogenetic Comparative Methods: A Meta-Analysis and Simulation Study. PLOS ONE 8:e67001.

Kubo, T., and Y. Iwasa. 1995. Inferting the Rates of Branching and Extinction from Molecular Phylogenies. Evolution 49:694–704.

Louca, S., and M. W. Pennell. 2020. Extant timetrees are consistent with a myriad of diversification histories. Nature 580:502–505.

Louca, S., and M. W. Pennell. 2021. Why extinction estimates from extant phylogenies are so often zero. Current Biology 31:3168–3173.e4.

Meyer, K., and W. G. Hill. 1992. Approximation of sampling variances and confidence intervals for maximum likelihood estimates of variance components. Journal of Animal Breeding and Genetics 109:264–280.

Morlon, H., S. Robin, and F. Hartig. 2022. Studying speciation and extinction dynamics from phylogenies: addressing identifiability issues. Trends in Ecology & Evolution 37:497–506.

Murphy, S. A., and A. W. Van Der Vaart. 2000. On Profile Likelihood. Journal of the American Statistical Association 95:449–465.

Neale, M. C., and M. B. Miller. 1997. The Use of Likelihood-Based Confidence Intervals in Genetic Models. Behavior Genetics 27:113–120.

Nee, S., E. C. Holmes, R. M. May, and P. H. Harvey. 1994. Extinction rates can be estimated from molecular phylogenies. Phil. Trans. R. Soc. Lond. B 344:77–82.

R Core Team (2022). R: A language and environment for statistical computing. R Foundation for Statistical Computing, Vienna, Austria. URL https://www.R-project.org/.

Rabosky, D. L. 2010. Extinction Rates Should Not Be Estimated from Molecular Phylogenies. Evolution 64:1816–1824.

Stadler, T. 2013. How Can We Improve Accuracy of Macroevolutionary Rate Estimates? Systematic Biology 62:321–329.

Tsimring, L. S. 2014. Noise in Biology. Reports on progress in physics. Physical Society (Great Britain) 77:026601.

Uyeda, J. C., and L. J. Harmon. 2014. A Novel Bayesian Method for Inferring and Interpreting the Dynamics of Adaptive Landscapes from Phylogenetic Comparative Data. Systematic Biology 63:902–918.

Venzon, D. J., and S. H. Moolgavkar. 1988. A Method for Computing Profile-Likelihood-Based Confidence Intervals. Journal of the Royal Statistical Society: Series C (Applied Statistics) 37:87–94.

Wieland, F.-G., A. L. Hauber, M. Rosenblatt, C. Tönsing, and J. Timmer. 2021. On structural and practical identifiability. Current Opinion in Systems Biology 25:60–69.

